# The Anaphase Promoting Complex targets the toxic protein Progerin for ubiquitin-dependent degradation via autophagy

**DOI:** 10.64898/2026.07.23.740245

**Authors:** Valeria Martinez, Matthew Lubachowski, Zoe E. Gillespie, Morgan Fleming, Troy A. A. Harkness, Christopher H. Eskiw

**Author notes:** Co-corresponding authors: Troy A. A. Harkness,; Christopher H. Eskiw. Co-first authors (Valeria Martinez, Matthew Lubachowski). Email list: Valeria Martinez, Matthew Lubachowski, Zoe E. Gillespie, Morgan Fleming.

## Abstract

The premature aging disease Hutchinson-Gilford Progeria Syndrome (HGPS) results from the accumulation of progerin, a cytotoxic protein generated from a point mutation in the *Lamin A/C* gene, in the nuclear lamina. Upon the proper stimulation, cells degrade progerin, reversing cellular HGPS phenotypes; however, there is still a gap in our knowledge concerning which pathways are mediating progerin degradation. Previous data has demonstrated that the Anaphase Promoting Complex (APC), a multi-subunit ubiquitin ligase, tagets proteins for degradation, and that a decrease in APC function is linked with cellular aging. To determine if the APC is linked to HGPS disease phenotypes, we performed a meta-analysis of RNA-seq data from skin samples isolated from HGPS patients and identified dysregulation of several genes encoding subunits and substrates of the APC. Stimulation of APC activity decreased progerin protein levels and significantly decreased the number of cells with nuclear blebs. Proximity ligation assays (PLA) demonstrated that APC structure is compromised in HGPS cells and that APC stimulation increases proximity of the APC with progerin. Coimmunoprecipitation revealed that the APC co-activator, CDC20, physically interacted with nuclear lamina proteins. We further demonstrate that APC-mediated progerin degradation occurs through autophagy. Inhibition of the 26S proteasome enhanced progerin degradation, providing additional support for APC mediated-progerin degradation occurring independent of the proteasome. As such, we propose a previously unidentified interaction and mechanism by which cells remove progerin. This finding has impact on potential therapeutic strategies for HGPS, as well as providing further insight into linking the APC with both normal and premature aging.

## INTRODUCTION

Hutchinson Gilford Progeria Syndrome (HGPS) is a rare childhood disease resulting from a mutation in the *LaminA* gene and the expression of the cytotoxic protein, progerin (1–3). This mutation results in the activation of a cryptic splice site (4, 5), generating an internal deletion of 50 amino acids, which includes a cleavage site for the zinc metallopeptidase STE24 (ZMPSTE24) (6, 7). The deletion of this site prevents farnesylated progerin from being cleaved and liberated from the lamina in patient cells, resulting in numerous cellular defects, including the loss of heterochromatin and dysregulation of gene expression (2, 7–11), as well as striking deformation of the nuclear membrane and lamina (6). There are currently no cures for this disease and, although some of the treatments, such as farnesyl transferase inhibitors (FTIs; (6, 12, 13), increase lifespan, this is only by a few years and with significant side effects that also impact quality of life.

Given the severity of HGPS and it’s theorized mechanistic links to normal aging, research has been dedicated to slowing or even curing this disease. Recent publications demonstrate a potential for the use of genetic modification and CRISPR approaches to eliminate progerin expression but viable treatments are decades away and would require embryonic screening with, potentially, germ-line editing. Although current treatments target the modification of progerin with farnesyl groups with the aim of preventing accumulation in the nuclear lamina, this inhibits the farnysaltion of other proteins, such as the RAS oncoprotein. These treatments also demonstrate limited impact on lifespan with poor quality of life outcomes. An alternative to this that can benefit patients already living with HGPS is to promote the removal or degradation of progerin at the cellular level. This has shown promise, with the inhibition of the mammalian target of Rapamycin complex I (mTORC1) with various pharmacological agents promoting reduction of progerin levels (14–18). It has been shown that reduction in progerin levels is through ubiquitin-mediated autophagy (16, 19), which is further stimulated by inhibition of other protein degradation machinery, such as the 26S proteasome (20, 21). Furthermore, modulation of other cellular energy sensing pathways with compounds, such as metformin (22–24), also promote autophagy and progerin degradation. Although this has also shown promise, clinical applications using these compounds has not materialized; however, these data do indicate important autophagic/cell maintenance mechanisms involved in progerin degradation and promotion of proper cellular function. As such as, investigating mechanisms that can be stimulated to promote progerin degradation would be of significant interest.

Ubiquitin is a small peptide (8.5 kDa) that, when tagged to proteins, targets them for degradation. There are a myriad of enzymes, called E3 ubiquitin ligases that are responsible for conjugating ubiquitin to target proteins. The Anaphase Promoting Complex (APC) is a major multi-subunit, evolutionarily conserved E3 responsible for protein degradation tied to cell cycle progression through mitosis and G1 (25–27). The APC is controlled by 2 mutually exclusive co-activators: CDC20, which drives entrance to mitosis, and CDH1, required for mitotic exit and G1 maintenance. It has become clear over the last decade that mis-regulation of the APC plays important roles in cancer progression, aging, and stress response (28–32). Consistent with a role for the APC in maintaining genome stability and a healthy lifespan, mutations to APC subunits and the APC co-activator FZY1/CDH1 in mammalian cells results in genome instability, cancer progression, and premature replicative aging (26, 33, 34). The APC targets many oncogenic kinases and transcription factors that promote proliferation for degradation, while maintaining levels of proteins involved in stress management and hormesis. As such, APC impairment leads to increased oncogenic potential, with APC activation resetting this imbalance (26). Therefore, the APC is potentially poised to act in a positive manner in HGPS cells by targeting proteins that inhibit the stress response for degradation, thereby reversing HGPS phenotypes.

Importantly for cell cycle regulation, the APC is inhibited by the Spindle Assembly Checkpoint (SAC) prior to mitosis to ensure all chromosomes are properly aligned along the metaphase plate prior to division (35). A recent study found that multiple SAC components were downregulated at the transcriptional and protein level in skin cells expressing progerin and in HGPS patient fibroblasts (36). A set of mouse studies demonstrated that low expression of the SAC component BUBR1 resulted in progeroid-like phenotypes (37) and that BUB3/RAE1 happlo-insufficuient mice also show signs of premature aging and a progeroid-like state (38). Since the SAC inhibits the APC, the downregulation of SAC components suggests that the APC may be more active in HGPS cells.

Once the chromosomes are properly attached to the mitotic spindle and are aligned along the metaphase plate, CDC20 dissociates from the SAC component MAD2, and is recruited to the APC, promoting mitosis (35). The small chemical molecule, Mad2-inhibitor-1 (M2I-1), inhibits the interaction between CDC20 and MAD2, maintaining higher than normal levels of APC activity, resulting in increased degradation of APC substrates (39). This has implications in protein turnover within cells, promoting proetostasis by enhancing the degradation of cellular proteins that can accumulate and damage cells. The use of M2I-1 has been shown to increase lifespan in yeast cells (40), as well as resensitize multiple drug resistant cancer cells to therapeutics (26, 41, 42).

Given the importance of the APC in aging, and evidence that progerin degradation reverses HGPS cellular phenotypes, we performed a meta analysis of published RNA-seq data generated from HGPS cells (43) to determine if the APC function was impacted in HGPS. In addition to many other genes and pathways, we identified several APC substrates and subunits that were dysregulated at the transcript level. Furthermore, global analysis of genes changing expression in HGPS exhibited enrichment of pathways associated with the APC, leading us to hypothesize that APC function is altered in HGPS cells and that APC activation in HGPS cells may contribute to phenotype reversal. To test this, HGPS cells were treated with M2I-1 to up-regulated APC actitivty, whichidentified decreased protein levels of progerin and lamin A/C. This decrease is concordant with a decrease in the number of cells exhibiting nuclear blebs, highlighting that the APC may also play a role in mediating progerin turn over. Proximity ligation assays (PLA) demonstrated that interactions between APC subunits CDC27 and APC2 were infrequent in HGPS cells and that treatment with M2I-1 significantly increased detectable interactions, as well as promoted interactions between APC subunits with progerin. PLA confirmed increased PLA proximity between ubiquitin and progerin in response to M2I-1. Inhibition of the 26S proteosome increased levels of progerin degradation while inhibiting autophagy prevented progerin degradation. These data demonstrate a previously unidentified interaction between the APC and progerin, indicating that autophagy is the primary mechanism by which APC-mediated ubiquitination promotes progerin clearance.

## Materials and Methods

### Cell culture

This study used the following cell lines: 1) immortalized fibroblasts (NB1-hTERT); 2) NB1-hTERT fibroblasts stably expressing GFP-progerin; and 3) human primary dermal fibroblasts isolated from skin biopsies of HGPS patients (HGADFN167 and HGADFN169, from the Progeria Research Foundation). HGADFN167 cells are derived from a male at 8 years and 5 months, while the HGADFN169 cell line is from a male at 8 years and 6 months (Progeria Research Foundation, ExPASY Bioinformatics Resource Portal). Fibroblasts were cultured in 1X Dulbeccos’s Modified Eagle Medium (DMEM; pH 7.7, Corning, USA, Cat# ca45000-304), containing high glucose (4.5 mg/mL), sodium pyruvate, and glutamine, supplemented with fetal bovine serum (FBS; 10% for NB1-hTERT cells and 20% for progeria cells; Gibco, Thermo Fisher Scientific, USA, Cat#: 12483-020) and 1.0% penicillin-streptomycin (GE Healthcare Life Sciences, USA, Cat#: SV30010). Cultures were maintained under standard culture conditions: 37°C with 5% CO_2_ in a humidity-controlled incubator. All cells were plated at 3,000 cells/cm^2^ for normal fibroblasts and 4,500 cells/cm^2^ for HGPS fibroblasts in either dishes or on 22 cm^2^ glass coverslips for imaging applications. The media was changed every 3-4 days.

### Cell passage

Cells were passaged when they reached ∼70% density to avoid contact inhibition. Cells were incubated with 3 ml of Tryple Express (Life Technologies, USA, Cat#: 12604013) for 3 to 5 minutes, or until cells were entirely disassociated from the plate surface. Using 7 mL of media, the cells were rinsed from the plate and collected into a 15 mL conical tube, followed by centrifugation at 300*g* for 5 minutes (Eppendorf, Hamburg, Germany). Supernatant was discarded, and the resulting pellet was resuspended in 10 mL of media. 10 µL of the suspension was mixed with 10 µL of 0.4% Trypan Blue Solution (Life Technologies, Canada, Cat# 15250061). The cell concentration was calculated using an automated cell counter (Countess^TM^ -Invitrogen).

To determine the population doubling times (PDT), the following equation was used:

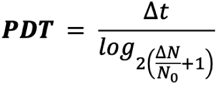

Δ*N* is the difference between the final number of cells and the initial count, representing the number of cells grown during the time interval Δ*t*.

### Cell treatments

All treatment compounds, except for MG132 (Sigma, Canada, Cat#1211877-36-9), were reconstituted in dimethyl sulfoxide (DMSO). To induce APC activity, the APC activator drug, Mad2 Inhibitor 1 (M2I-1; (39) was evaluated at concentrations ranging from 1-20 µM for 24-72 h. To inhibit the APC, we used APCIN (44) Thermo Scientific, Cat#50-205-3972) at concentrations ranging from 5-10 µM. Additionally, MG132 was used at concentrations of 5-10 µM to inhibit the proteasome. Before use, MG132 was reconstituted in molecular-grade water and filtered sterilized. Chloroquine (Sigma Aldrich, cat# 50-63-5) was used at a final concentration of 5 µM in culture media for a 48-h period, either alone, or in combination with 5 µM of M2I-1. Western blot analyses and densitometry were used to evaluate protein abundance, as well as colocalization analyses of cells grown on 22 mm^2^ coverslips. All measurements and assays were performed on 3 biological replicates to ensure the validity and reproducibility of the data/observations.

### Western Blotting (WB), Immunoprecipitation (IP), and Immunofluorescence (IF)

WB, IP and IF were was performed according to previously published protocols (41, 45). The following primary antibodies were used in a 1:1000 dilution: mouse anti progerin (Santa Cruz, cat#SC-81611), mouse anti Lamin A/C (Santa Cruz, cat#sc-376248) and rabbit anti cyclin B1 (Sigma, C8831), mouse anti Lamin A/C (Santa Cruz, cat# sc-376248), rabbit anti CDC20 (Sigma, cat#, sc-5296). The secondary antibodies used were goat anti rabbit HRP (dilution 1:2500, Jackson Scientific, cat#, 115-035-003), and goat anti mouse HRP (dilution 1:1500, Jackson Scientific115-035-144).

### Proximity ligation assay (PLA)

The PLA reaction was performed using Duolink® PLA from Sigma (Cat #: DUO92202-1KT). To assess the ideal concentration of the antibodies for use in PLA, the performance of the antibodies was evaluated in individual IF reactions. After visualization of the images, the antibody concentrations selected were 1:200 for APC2, 1:50 for CDC20, 1:50 for progerin, 1:50 for CDC27, and 1:50 for ubiquitin. The antibody combinations used were progerin-APC2, CDC27-APC2, CDC20-APC2, CDC20-progerin and ubiquitin-progerin. The PLA reactions were performed according to the manufacturer’s instructions. Briefly, cells were grown on 1 cm^2^ sterilized cover slides to 70% confluency, then fixed and permeabilized, as described above. Next, 1 drop of Duolink® Blocking solution was added to each sample. The samples were incubated cell-side down on parafilm in a heated humidity chamber for 60 min at 37°C. Next, the diluted primary antibodies were added and incubated for a further 1h at 21°C, washed in buffer A (prepared according to the manufacturer’s instructions), then incubated with the PLUS and MINUS PLA probes for 1 h at 37°C. After 2 washes with buffer A, the samples were incubated with ligase for 30 min at 37°C and then with polymerase, diluted in the appropriate buffer, as recommended by the manufacturer, for amplification. The amplification was performed at 37°C for 100 min. After final washes in buffer B, the slides were mounted in VectaShield^TM^ with DAPI. The cells were visualized using a Ziess X61 microscope; gain and exposure times were optimized for each channel, and the number of foci of cells per treatment were quantified. A minimum of 30 nuclei were counted for each condition.

### Colocalization

HGPS primary fibroblasts were treated with or without 5 µM of chloroquine in combination with the vehicle control DMSO or the APC activator M2I-1. Double protein staining was performed by incubating the cells simultaneously with primary antibodies against mouse progerin (dilution 1:200, Santa Cruz; Cat #; SC-81611) and rabbit ubiquitin (Proteintech; Cat #: 10201-2-AP). This was followed with the secondary antibodies Alexa Fluor 488 anti-mouse and donkey anti-rabbit cy3 (Jackson ImmunoResearch; Cat #: 711-165-152) as detailed above. Images of cells were collected at 100x magnification with constant exposure times. Gray-scale images were imported into Photoshop and converted to RGB files.

### NB1-hTERT GFP-progerin stable cell line

NB1-hTERT fibroblasts growing in 6-well dishes (∼80% of confluency) produced a non-clonal population that stably expressed GFP-progerin or GFP. For transfection, NB1-hTERT fibroblasts were incubated for 3 h in Opti-MEM (ThermoFisher Scientific; Cat #: 31985062). Transfections were carried out using lipofectamine™3000 (ThermoFisher Scientific; Cat #: L3000001) according to the manufacturer’s instructions. Briefly, 5 µg of DNA (linearized plasmid), 5 µL of P3000™, and 7.5 µL of the Lipofectamine™3000 reagent were added to the cells, which were then incubated for 6 h. Next, 2 mL of regular media were added to each well (DMEM with 10% FBS and 1% P/S Opti-MEM), followed by a 24 hr incubation. Media was replaced by selection media consisting of 500 µg/mL of G418 sulfate (Thermo Fisher Scientific, USA: Cat #: 11811031) in DMEM. Cell growth and survival was monitored visually. Cells were fixed and monitored to confirm plasmid insertion by immunofluorescence and western blot.

### Nuclear fractionation

NB1-hTERT immortalized fibroblasts expressing GFP-progerin were washed in PBS and harvested. Cells were pelleted by brief centrifugation for 10 s at 8600 x g, and resuspended in PBS containing 0.1 % NP-40. The cell suspension was triturated five times through the tip of a P1000 micropipette, and a portion was aliquoted to serve as the whole cell fraction. The remaining lysate was again centrifuged as above. A portion of the supernatant was reserved as the cytoplasmic fraction, with the remainder discarded, and the pellet was washed via resuspension in PBS containing 0.1% NP-40 followed by another round of centrifugation and resuspension to become the nuclear fraction. All fractions had Laemmli buffer added to 1X, and the whole cell and nuclear fractions were briefly sonicated for 5 s on ice with a 30% duty cycle. All samples were boiled for 5 minutes then stored at −20°C until analysis.

### RNA-seq Meta-Analyses

Data used in metaanalyses were retrieved from GEO record GSE113957 (43). All HGPS RNA-seq datasets were retrieved (n=10), and all appropriately age-matched controls within range of HGPS dataset ages (n=18) (Table S1). Raw RNAseq reads were examined for quality using FastQC (https://www.bioinformatics.babraham.ac.uk/projects/fastqc/). TrimGalore! was used to complete trimming, removing adaptors and poor quality reads (https://github.com/FelixKrueger/TrimGalore). Data were then aligned using Hisat2 (46) to GRCh38 and resultant bam files imported into SeqMonk (Babraham Bioinformatics, Cambridge, UK). HGPS and age-matched controls were grouped and differential gene expression analyses were completed in SeqMonk, using the built in RNA-seq quantiation pipeline (around the gene, raw counts, single stranded) followed by DESeq2 (47). GO enrichment was completed using all significantly differentially expressed genes (false discovery rate < 0.05) using g:profiler! (48).

### Statistics

PLA analysis (control, M2I-1, or APCIN): 30 cells were chosen randomly, with the number of foci or PLA fluorescent dots counted. ImageJ was used to manually count the number of foci in each cell. A Kruskal test was used to examine differences in each treatment, followed by a Wilcox test to establish significant differences between the controls and treatments applied. Data was graphed in a box plot with error bars. For western blot analyses, data is presented as mean ± standard error of the mean (SEM). The significant differences between western band intensities was determined using a Student’s T-test. Fluorescence signal intensity was assessed after imaging at least 50 cells/treatment. Results were analyzed using the Kruskal Wallis test followed by a Pairwise Wilcoxon test with a Bonferroni correction of *p*-value, with outliers removed using interquantile range in R (49).

## Results

### APC substrate and regulatory RNAs are deregulated in HGPS fibroblasts

Given that the APC has been linked to cellular health and lifespan by regulating the turn over of proteins, we hypothesized that that the presence of progein in HGPS cells may also interfere with APC further leading to premature aging. To better understand the consequence of progerin expression on transcript profiles, we conducted a meta-analysis of RNAseq from HGPS patientfibroblasts to normal primary dermal fibroblasts (43). The datasets analysed included 3 male HGPS patients, 7 female HGPS patients and 18 from age-matched normal individuals (**Table S1**). Using a Principal Component Analysis (PCA), we observed that the normal age-matched datasets were clustered together, while the male and female HGPS datasets were more varied and distinct from age-matched controls (**Supplemental Figure 1**). This was reflected in differential gene analyses, identifying 1066 genes as significantly changing expression (525 up, 541 down; absolute fold change ≥ 2, FDR < 0.05) (**Tables S2-4**). Gene ontology (GO) analyses of genes significantly changing expression in HGPS (FDR < 0.05) revealed enrichment of APC-associated pathways, including ubiquitin-dependent protein catabolic process as well as ubiquitin ligase complex and cullin-RING ubiquitin ligase complex (both of which belong to the parent GO term anaphase-promoting complex; **Table S5**). Furthermore, we observed that many genes related to APC activity were either up- or down-deregulated in HGPS fibroblasts, compared to the normal controls (**Table S6**). The proteins encoded by these genes were either APC subunits, APC substrates, or involved with APC regulation. Several APC subunit/co-activator genes (*CDH1/FZR1*, *APC5*, and *APC2*) were slightly decreased, while two (*CDC16* and *APC16*) were elevated. Combined, these data provide evidence forAPC dysregulation in HGPS patient fibroblasts. It has been observed that the SAC is impaired in HGPS cells, suggesting that the APC may have increased activity (50). Furthermore, a study showing that mice expressing low levels of the SAC component BUBR1, or haploinsufficuent for both BUB3 and RAE1, expressed progeroid-like phenotypes (37) (38). Finally, inhibition of ubiquitination results in progerin degrradation in HGPS fibroblasts (21). Together, these previous observations as well as meta-analyses of RNAseq datasets, adds evidence to support the hypothesis that the APC is misregulated in progeria cells.

### APC activation decreases progerin protein levels

Our observations that APC-associated genes were deregulated in HGPS cells (HGADFN169) is an indication that the APC might play a role in progerin turnover in HGPS cells. As such, we focused on establishing a functional relationship between the APC and progerin-dependent cellular defects leading to premature aging. We first tested if APC substrates accumulate in HGPS cells. Western blotting of HURP, Cyclin B1 and CDC20 demonstrates that these APC subunits are present in higher levels in HGPS dermal fibroblasts than in normal dermal fibroblast cells (**Fig 1A**; 3 independent replicates shown). Progerin accumulation is shown in the HGPS cells while none is detected in normal fibroblasts, demonstrating that APC substrate accumulation occurs concomitantly with the presence of progerin.

**Figure 1.**
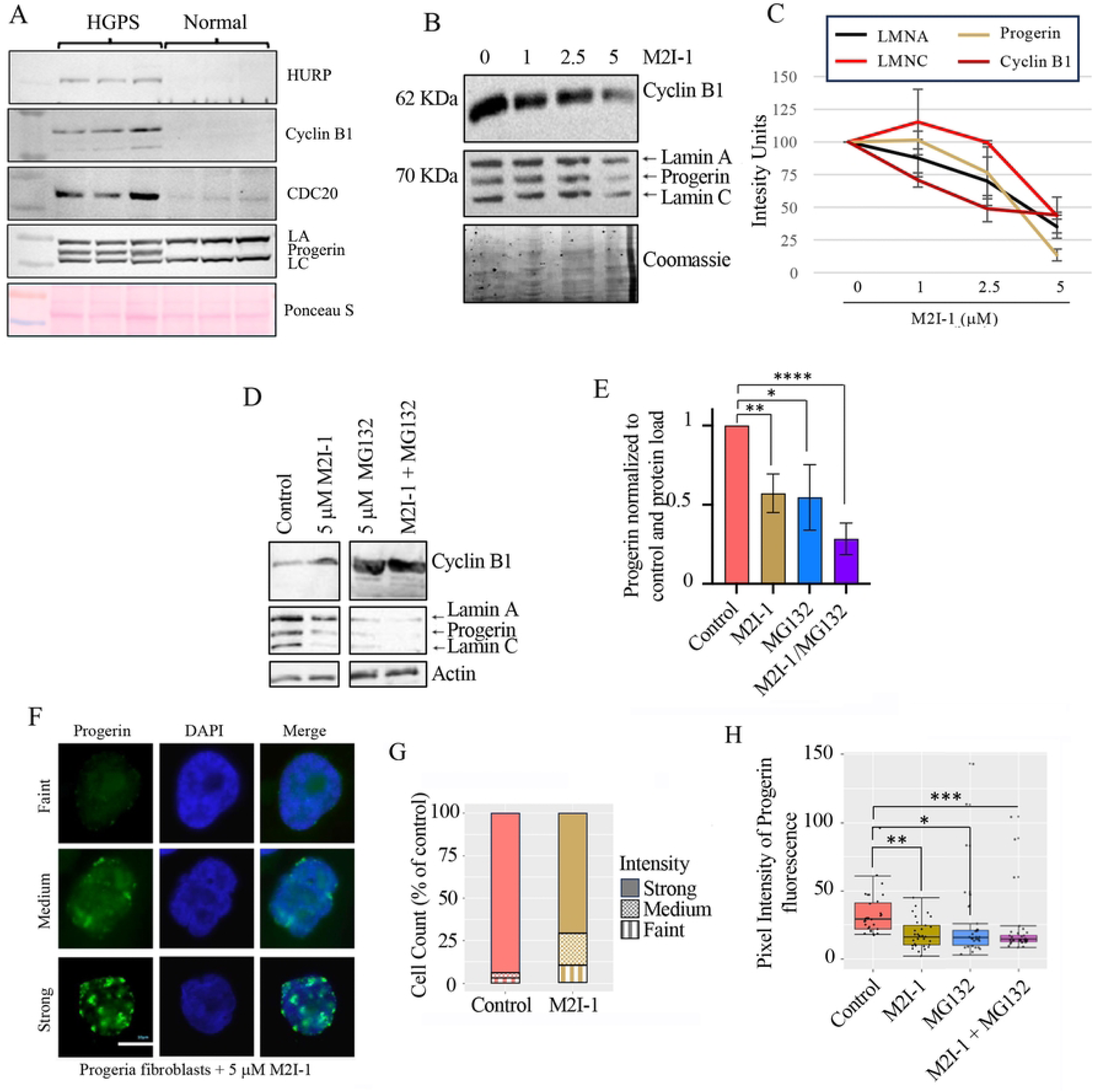
Activation of the APC reduces progerin protein levels in HGPS and Progerin expressing cells. A) Protein lysates from 3 independent cultures of HGADFN169 HGPS fibroblasts (HGPS) or normal dermal fibroblasts (Normal) were generated and western blotting performed for APC substrates, HURP, (1^st^ row), Clucin B1 (2^nd^ row) and CDC20 (3^rd^ row). Western blotting for Lamin A (LA), progerin (Progerin) and lamin C (LC) was also performed with load controls demonstrated by Ponceau S staining. B) HGADFN169 HGPS fibroblasts were grown in the presence of increasing concentrations of M2I-1 for 72 hours. Lysates were prepared with Westerns performed using antibodies against Cyclin B1 and Lamin A/C. Coomassie stained gels were used for load controls. C) Bands from two experiments as described in A) were quantified, averaged, normalized to the control and the Coomassie stained gel, then plotted. Arbitrary intensity units from the blots are plated on the Y axis. The SEM is shown. D) HGADFN169 cells were grown in the presence of 5 mM M2I-1 +/− 5 mM MG132, or left untreated, for 72 hours. Lysates were prepared and analyzed using antibodies against Lamin A/C and Cyclin B1. Actin was used to control for load. E) Quantification of the results from panel C was performed with all progerin bands quantified and plotted. n = 3, SEM is shown; * *p*-value < 0.5; ** *p*-value < 0.05; **** *p*-value < 0.0005. F) Progerin intensity levels in HGADFN169 cells growing on coverslips treated with 5 mM M2I-1 was determined by IF using antibodies against progerin. The levels observed were scored as faint, medium or strong. G) The percentage of HGADFN169 cells exhibiting strong, medium or faint intensity when labelled with antibodies against Progerin in the absence or presence of 5 mM M2I-1. H) The fluorescence intensity signal levels from ≥50 HGADFN169 cells/treatment were scored, with box plots presented. SEM is shown; * *p*-value = 0.0068; ** *p*-value = 0.00047; *** *p*-value = 2.4×10^-6^.

To test if APC activation has an impact on progerin levels in HGPS fibroblasts. HGPS fibroblasts (HGADFN169) were grown in the presence or absence of the small chemical APC activator, Mad2 Inhibitor-1 (M2I-1; (39)), for 72 hours. Western blotting of protein lysates was performed using antibodies against Lamin A/C andprogerin. We observed that progerin levels, as well as Lamin A and C, were reduced following M2I-1 treatment in a dose-dependent manner (**Fig 1B-1C**). Clycin B is a *bonafide* target of the APC with degradation of Cyclin B1 used as an indicator of APC activity (41). As such we used this as a control for evaluating the ability of M2I-1 to stimulate APC activity in HGPS cells. We observed Cyclin B1 was reduced by M2I-1 treatment, indicating that M2I-1 activates APC activity in HGPS cell. This is consistent with previously published data demonstrating M2I-1 reduces APC substrate protein levels in other systems (41, 42), indicating that the effect on progerin is specific to APC activation. We next tested if progerin turnover in response to M2I-1 is proteasome-dependent, as the APC targets many of its protein substrates for proteasome-dependent degradation. For this, HGPS fibroblasts were grown with M2I-1, in the presence or absence of the proteasome inhibitor MG132, for 72 hours. If the proteasome is required for progerin degradation in response to APC activation, we predicted that MG132 would abrogate this response. We observed that proteasome inhibition using MG132 also decreased progerin and Lamin A/C protein levels similar to those observed with M2I-1 (blots presented in **Fig 1D** and the quantification in **Fig 1E**). This is consistent with a previous report demonstrating that treating HGPS cells with MG132 leads to decreased progerin levels that was at least partially mediated by autophagy (20). However, when used in combination, M2I-1 and MG132 decreased progerin protein levels even further while Cyclin B levels accumulated, indicating that the APC did not target progerin degradation via the proteasome. This result indicates that the APC targets Cyclin B1 for degradation via the proteasome in HGPS cells, but progerin degradation is not impacted by the proteasome.

To confirm that progerin levels were reduced in M2I-1 treated HGPS HGADFN169 fibroblasts, we used indirect immunofluorescence to visualize progerin (**Fig 1F**). We measured pixel intensity from a minimum of 30 randomly selected cells, which was quantified and plotted, noting that strong progerin intensity was reduced in the presence of M2I-1. Flourescence intensity derived from progerin foci was reduced when cells were treated with M2I-1 (**Fig 1G**). We further quantified the intensity of progerin signal from untreated cells, or cells treated with M2I-1, MG132, and M2I-1 in combination with MG132, to evaluate the impact on progerin levels. We observed that M2I-1 decreased progerin signal intensity in HGADFN169 cells (**Fig 1H**). MG132 treatment also decreased levels of progerin, indicating that blocking the 26S proteosome stimulated cells to degrade this protein. M2I-1 in combination with MG132 also resulted in decreased progerin intensities compared to untreated controls. Our analysis is consistent with the hypothesis that the APC targets progerin for degradation independent of the proteasome.

### Activation of the APC reduces progerin phenotypes

If APC activation does reduce progerin levels, then it is expected that APC activation will impact HGPS phenotypes. We first assessed if treatment of HGPS fibroblasts with M2I-1 resulted in decreased number of cells exhibiting nuclear blebbing, a hallmark observed in HGPS cells (8). Nuclei in cells were counted and scored for whether they exhibited normal shape or nuclear abnormalities (**Fig 2A**). Nuclear blebbing defects were reduced in M2I-1 treated HGPS fibroblasts compared to untreated control cells (**Fig 2B**). We observed that 87.5% of control HGPS cells exhibited nuclear blebbing, while only 62.5% of M2I-1 treated cells showed this phenotype. We then asked whether APC activation impacted proliferation of HGPS cells. Previous observations showed that normal fibroblasts treated with Rapamycin exhibited reduced proliferation rates without induction of senescence (51). We treated two independent HGPS fibroblast lines (HGADFN167 and HGADFN169) with M2I-1, with cell doubling times and total cell doublings measured (**Fig 2C-2F**). We observed that cell doubling times in both M2I-1 treated cell lines increased, with a corresponding decrease in total doublings without causing cell death (data not shown), as observed in normal cells with Rapamycin (52). Taken together, our results thus far indicate that the APC is impaired in HGPS fibroblasts and can be activated leading to a reduction in progerin protein levels that is proteasome-independent, resulting in reduced nuclear blebbing and increased doubling times.

**Figure 2.**
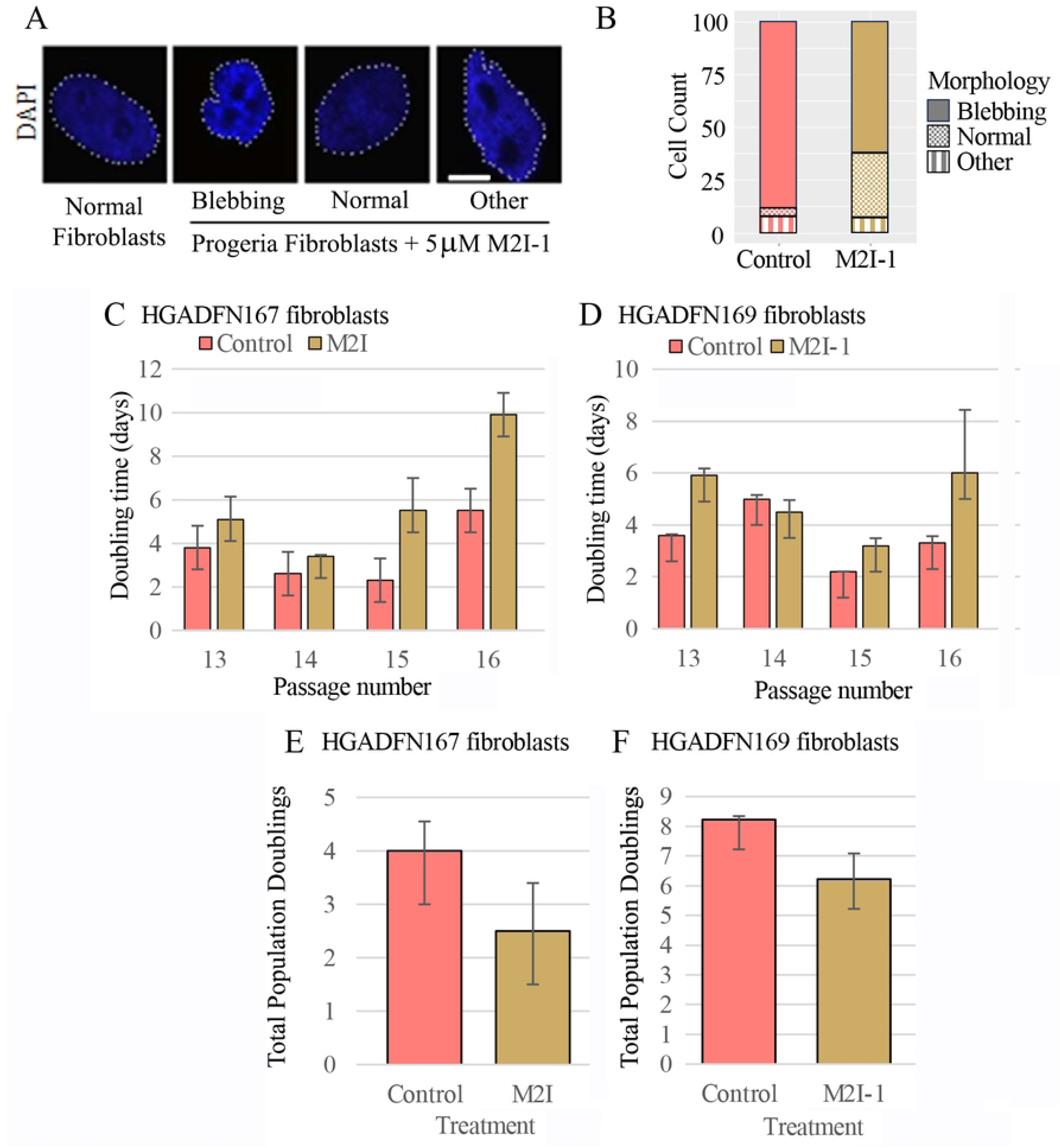
APC activation using M2I-1 improves nuclear membrane morphology and slows growth in HGPS fibroblasts. A) HGADFN169 HGPS fibroblasts were treated with 5 mM M2I-1 for 72 hours, then stained with DAPI to visualize nuclei. Images showing the different nuclear morphologies observed are presented. B) 100 cells were scored from control and M2I-1 treated cells for nuclear membrane morphology according to the scale shown in A). C) HGADFN167 HGPS fibroblasts growing in culture were counted every 4-5 days to determine the population doubling time (PDT). D) The PDT was determined for HGADFN169 HGPS fibroblasts. E) and F) The average number of cell doublings over 84 hours for M2I-1 treated and control HGADFN167 and HGADFN169 HGPS fibroblasts, respectively.

### Progerin interacts with CDC20 independent of the APC

We hypothesized that if the APC is targeting progerin for degradation, then the APC and progerin must be in close physical proximity, either directly interacting or within the same complex. To test this hypothesis, we used Proximity Ligation Assays (PLA), which indicates if two proteins are within close proximity (≤40 nm) to one another, as well as the localization of these interacting proteins within the nuclear volume (53, 54). As positive controls, we confirmed close proximity of the APC subunits APC2 and CDC27 in immortalized fibroblasts NB1-hTERT cells, which exhibited numerous foci (**Fig 3A – upper panels**) indicating that the reaction was able to detect proteins localizing within the APC complex. There were very few foci observed in HGADFN169 HGPS cells between APC2 and CDC27, indicating that APC structure may be altered (**Fig 3A – lower panels**). We repeated PLA using APC2 and CDC27 in HGPS cells treated with M2I-1 and APCIN (an inhibitor of APC function (44). The foci were counted from 30 cells for each treatment and plotted (**Fig 3B**; see **Suppl Fig 2** for representative images of nuclei; M2I-1 vs. Progeria control, *p*-value < 0.203; M2I-1 vs. APCIN, *p*-value < 0.024). We observed that M2I-1 increased the number of foci, demonstrating that M2I-1 promotes APC2-CDC27 interactions, whereas APCIN had no observable effect, as the number of foci remained low. M2I-1 releases the APC co-activator CDC20 from the SAC (39), allowing premature interactions with, and activation of, the APC. In order for CDC20 to bind the APC, CDC27 must be phosphorylated, which leads to the phosphorylation of APC1, and a conformational change that allows CDC20 to bind (50). We showed previously in MCF7 breast cancer cells that M2I-1 increased the content of phosphorylation of APC1^S355^ (41). We interpret the observation to mean that M2I-1 increases APC2-CDC27 interactions, indicating that M2I-1-dependent release of CDC20 from the SAC may lead to CDC27 phosphorylation, which in turn stimulate APC2-CDC27 interactions.

**Figure 3.**
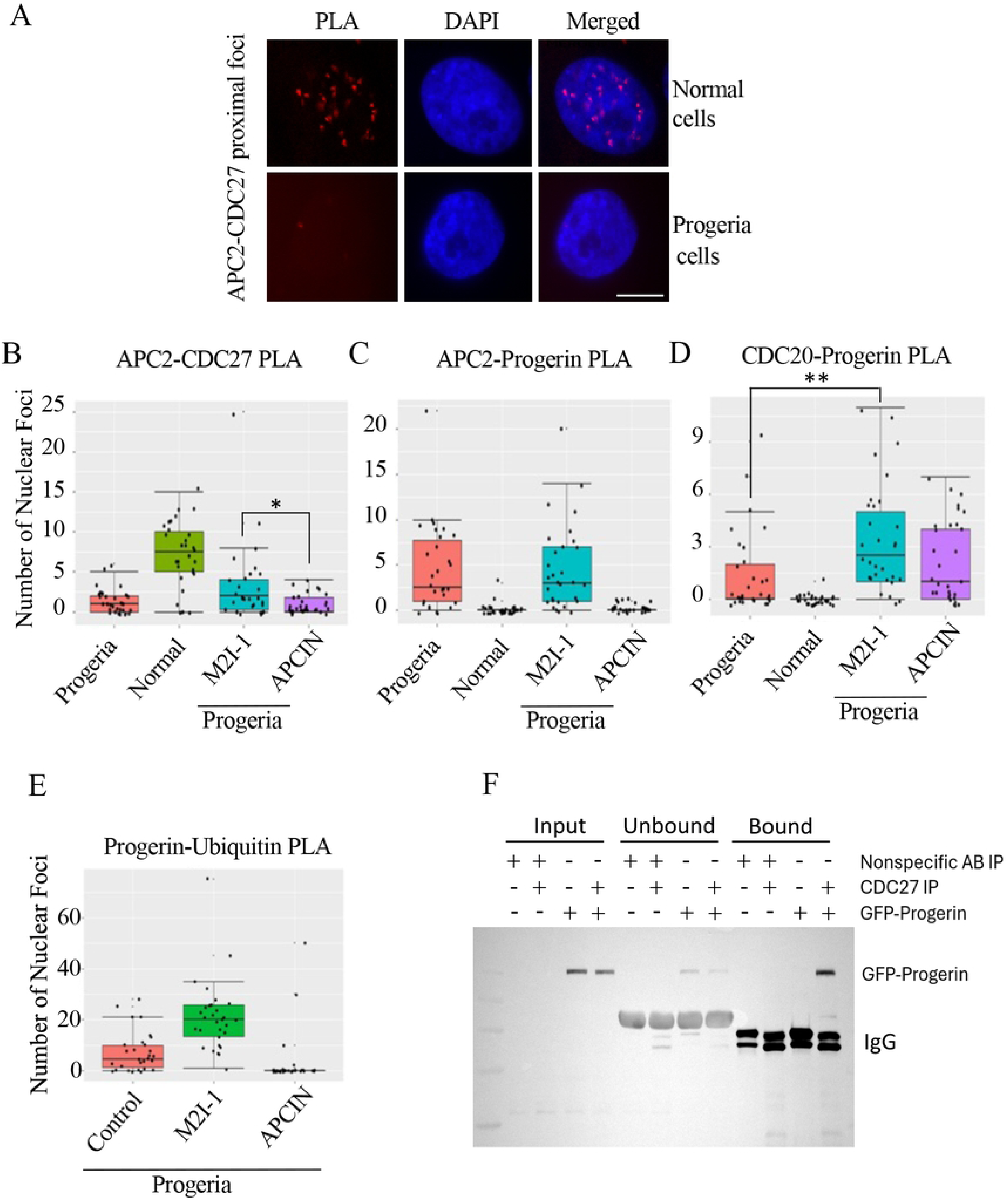
APC activation in HGADFN169 cells increases APC subunit proximity, and enhances the proximity of progerin with CDC20 and ubiquitin. A) The Proximity Ligation Assay (PLA) was used to determine APC subunits APC2 and CDC27 were proximal. Foci (red) within the nucleus (blue) indicate that the 2 proteins are within at least 40 nm. NB1-hTERT cells were used as a normal control. PLA was used to indicate proximity of proteins pairs: B) APC2-CDC27; C) APC2-Progerin; and D) CDC20-Progerin. Foci (red) from 30 nuclei were counted from each treatment and shown as a box plot with error bars. E) PLA was used to assess the proximity of progerin and ubiquitin. HGADF169 cells were treated with either DMSO, the APC activator M2I-1, or the APC inhibitor APCIN. The foci from 30 random nuclei were counted and shown as box plots with SEM shown. F) Lysates prepared from NB1-hTERT cells stably expressing GFP or GFP-Progerin were immuno-precipitated using antibodies against Lamin A/C, which recognize progerin. The immunoprecipitates were analyzed with antibodies against Lamin A/C and CDC20.

We next performed PLA against APC2 and progerin (**Fig 3C; see Supp Fig 3** for images of cells). In HGPS fibroblasts, there were foci between APC2 and progerin, indicating that progerin is in close proximity to the APC. In normal cells, there were no interactions observed, as expected since normal cells do not express detectable levels of progerin (**Fig 3C; see Supp Fig 4A**). However, M2I-1 treatment increased the number of foci observed between APC2 and progerin, but APCIN completely abolished the proximity of APC2 and progerin. APCIN inhibits the interaction between the CDC20 co-activator with the APC; therefore, our observations indicate that CDC20 is required for the interaction between APC2 and progerin. To test this hypothesis, we performed PLA using antibodies against CDC20 and progerin (**Fig 3D; see Supp Fig 4B and 5 for images**). As negative controls, we tested interactions between CDC20 and progerin in non-HGPS cells and observed no signal, indicating no observable background signal from PLA reactions (**Fig 3D and Supp Fig 4B**), and, as with APC2, we observed that CDC20 was in proximity to progerin in HGPS fibroblasts. M2I-1 produced a large increase in observed foci (*p-*value = 0.0042), indicating that freeing CDC20 from the SAC allows CDC20 to be within proximity of progerin. Of significance, we observed that APCIN did not inhibit this interaction, indicating that the interaction of CDC20 with progerin does not require the APC.

**Figure 4.**
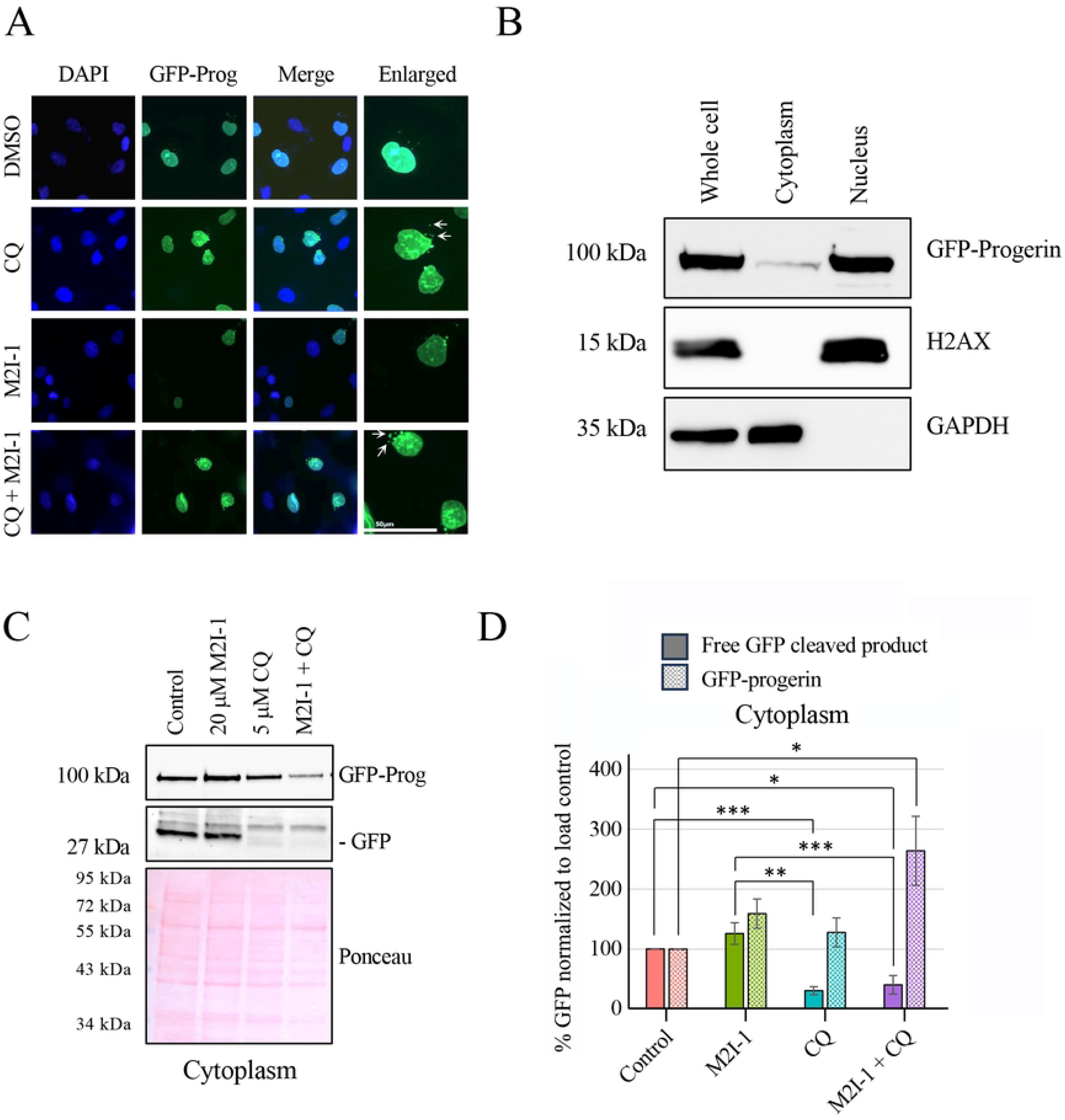
Inhibition of autophagy reduces APC-dependent degradation of progerin. A) NB1-hTERT GFP-progerin cells were treated for 48 h with carrier (DMSO), +/− 5 mM chloroquine (CQ) 5 mM M2I-1 (M2I-1) or chloroquine with 5 mM M2I-1 (CQ + M2I-1). Green: GFP-progerin, Blue: DAPI - chromatin. White arrows indicate the accumulations of GFP-progerin outside the nucleus in CQ treated cells. B) Whole cell, nuclear and cytoplasmic protein fractions were generated from NB1-hTERT GFP-progerin cells, were analyzed for levels H2AX (nucleus), GAPDH (cytoplasm) and GFP (GFP-progerin). C) NB1-hTERT GFP-progerin cells were either untreated (Control), treated with 20 mM M2I-1, 5mM chloroquine (5mM CQ), or 20 mM M2I-1 with 5mM chloroquine (M2I-1 + CQ). The cytoplasmic fractions were retained. Full length GFP-progerin and cleaved GFP resistant to autophagic breakdown were asseseed by Western blot. The Ponceau S stained membrane was used as the load control. D) Quantified images and values are displayed. Intensities were averaged, normalized to the control and the load control, then plotted. Solid bars represent free GFP cleaved products, with hatched bars presenting GFP-progerin. SEM is shown with n=5. * p-value < 0.05; ** p-value < 0.005; *** p-value < 0.0005.

**Figure 5.**
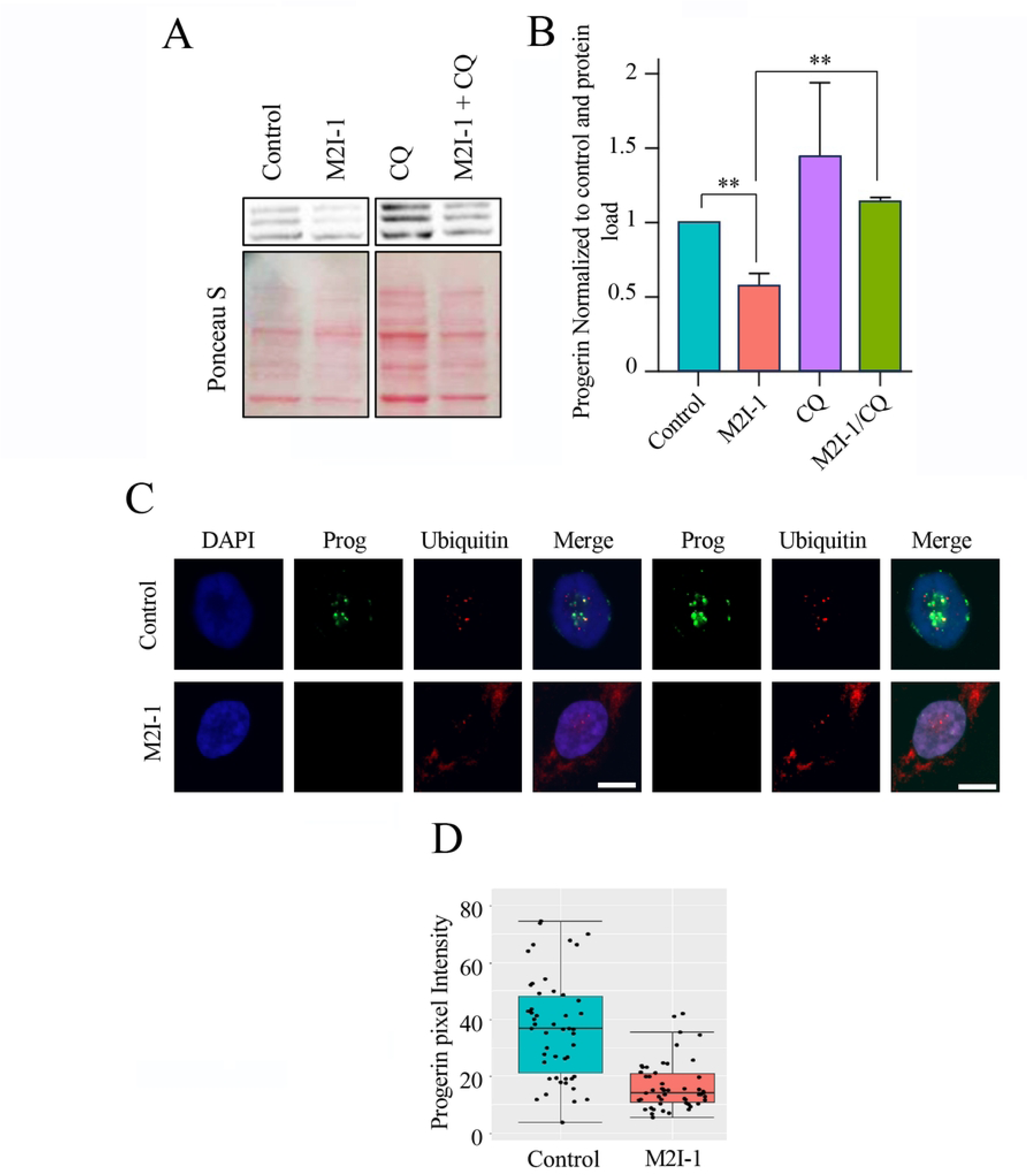
Inhibition of autophagy blocks turnover of ubiqutinated nuclear proteins. A) HGADFN169 HGPS fibroblasts were treated with 20 µM M2I-1 (M2I-1) or 5 µM chloroquine (CQ) or M2I-1 + chloroquine (M2I-1 + CQ). Cell lysates were prepared and analyzed with antibodies against Lamin A/C. Ponceau S stained gels were used as load controls. B) Three separate experiments were performed for each cell line, with all bands quantified, compared to load control, combined, and plotted. SEM is shown, **, p-value < .005. C) HGADFN169 HGPS fibroblasts grown on coverslips were treated with 20 µM M2I-1 (M2I-1). After 48 hours, cells were fixed, permeabilized and immunolabeled with for progerin or ubiquitin. Images were collected and false colored. The same images are presented but with the signal intensity increased. Although this saturates some pixels it also allows for the visualization of the suble signals that are otherwise missed. Green: progerin; red: ubiquitin; blue: DNA. D) Pixel intensity from at ≥50 cells was measured for each treatment and plotted as a box plot. Note that saturated images were not qualtified in this plot as this would distort the data.

To further test the hypothesis that CDC20 binds progerin to recruit it to the APC for ubiquitination, we performed an additional PLA using antibodies against progerin and ubiquitin (**Fig 3E; see Supp Fig 6 for images**) in HGADFN169 cells. We observed foci in the HGPS fibroblasts indicating that progerin is in proximity to ubiquitin. We treated cells with either M2I-1 or APCIN to assess the role that the APC plays in progerin ubiquitination. M2I-1 markedly increased the ubiquitin-progerin foci observed in HGPS cells. This is consistent with M2I-1 increasing CDC20-progerin foci in HGPS fibroblasts (**Fig 3D**). Strikingly, APCIN completely abolished progerin-ubiquitin foci in these cells. While APCIN does not inhibit the apparent interaction between CDC20 and progerin, it completely blocks its proximity to ubiquitin. To further validate our hypothesis that CDC20 and progerin are within proximity, we used antibodies against CDC27 to immunoprecipitate (IP) progerin in NB1-hTERT control cells, and in cells expressing GFP-progerin. We used GFP antibodies in Western blots to detect GFP-Progerin (**Fig 3F**). GFP-Progerin was observed when CoIPed with CDC27 but not with nonspecific control antibodies. In conclusion, our observations indicate that progerin interacts with the APC, via CDC20 first binding Progerin, and then delivering it within proximity of the APC in order to be ubiquitinated.

**Figure 6.**
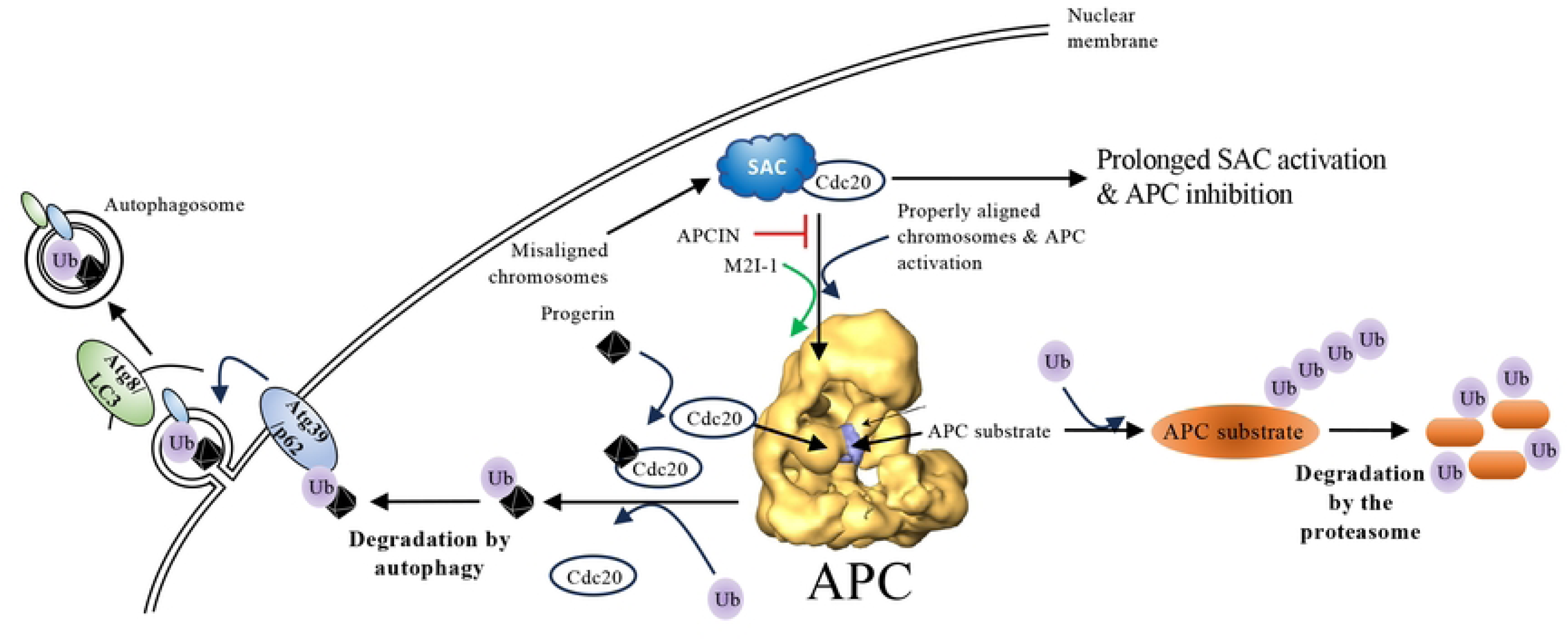
Proposed mechanism for APC activation and targeted autophagy of cytotoxic nuclear proteins. Cytotoxic protein aggregates (black diamond) are predicted to interact with CDC20 for recruitment to the APC, which are then ubiquitinated by the APC to enter the autophagic pathway. The APC normally targets cell cycle regulated proteins for proteasomal degradation. The conserved autophagy pathway members p62 and LC3-II are predicted to act in this pathway. Reduced APC activity promotes aging possibly by inhibiting autophagy. Experimental activation of the APC is through M2I-1 (disrupts SAC/CDC20 interactions), and APC inhibition is through APCIN (inhibits recruitment of CDC20 to the APC).

### Induction of progerin degradation is via APC-mediated autophagy

Our analyses indicated that progerin is targeted for ubiquitination and degradation by the APC and that APC activation in HGPS fibroblasts increases progerin turnover. However, unlike the canonical APC substrate Cyclin B, progerin is not targeted for degradation through the proteasome (**Fig 1C**). The other major protein degradation pathway utilized by cells is autophagy (55). Autophagy induction extends lifespan of most models tested (56) with rapamycin and other mTOR inhibitors (57), linked to reductions in progerin levels (16, 58). As such, we treated NB1-hTERT GFP-progerin cells, with 5 μM M2I-1 in the presence or absence of 5 mM chloroquine (CQ), which blocks autophagy (59) (60). We imaged cells and observed that GFP levels were low in M2I-1 treated cells, but higher when cells were treated with both M2I-1 and CQ (**Fig 4A**). We observed that chloroquine increased progerin fluorescence in control and M2I-1 treated cells, indicating that APC-targeted degradation of progerin utilizes autophagy, not the proteasome. Furthermore, cytoplasmic GFP foci were observed in the presence of CQ (**Fig 4A, white arrows**). Specifically, Over 300 cells were examined for the presence of cytoplasmic foci in each treatment. The percent of cells with cytoplasmic foci in control NB1-cells GFP-progerin was 4.9%. When M2I-1 was used, it was 5.5%, and when CQ was used, the total was 14.9%. When CQ and M2I-1 were both used, the total did not increase and remained at 14.4%. These data support the conclusion that progerin is targeted for APC-mediated autophagy.

To assess the role of autophagy in the degradation of progerin, we used NB1-hTERT GFP-progerin cells and Western blotting to determine whether increased GFP-progerin can be observed in the cytoplasm in the presence of M2I-1. We fractionated cells into nuclear and cytoplasmic fractions. GFP is relatively resistant to degradation in the lysosome (61); free GFP accumulates in the lysosome when a GFP-tagged protein, like GFP-progerin, is targeted for autophagy. Fractionation of the cells showed that the bulk of GFP-progerin is in the nucleus, with a small amount in the cytoplasm (**Fig 4B**). Lysates of the fractions were analyzed using antibodies against GAPDH and H2AX as markers of the cytosol and the nucleus, respectively. The cells were then treated with 20 μM M2I-1 +/− 5 μM CQ, and fractionated to produce cytoplasmic lysates. We observed that free GFP accumulated in the cytoplasm of M2I-1 treated cells (**Fig 4C**). CQ inhibited the cleavage of GFP when used alone or in combination with M2I-1, consistent with the APC targeting GFP-progerin for degradation via autophagy (**Fig 4C**). Quantitation of the bands from 5 repeats of the M2I-1 +/− CQ experiment, with normalization to untreated controls and the Ponceau S stained membrane, showed that CQ inhibited the cleavage of GFP-progerin in M2I-1 treated cells and that M2I-1 + CQ increased GFP-progerin levels in the cytoplasm (**Fig 4D**). This additionally supports the observations that M2I-1 induces degradation of GFP-progerin in an autophagy-dependent manner.

To provide further evidence that progerin is being degraded through an APC-dependant autophagy mechanism, we treated HGADFN169 HGPS fibroblasts with 5 μM M2I-1 +/− 5 μM CQ for 48 hours. Lysates were prepared and analyzed using antibodies against Lamin A/C (**Fig 5A**). Quantification of bands from each blot for each cell line, done in triplicate, was performed and combined (**Fig 5B**). Similar to observations made using NB1-hTERT GFP-progerin cells (**Fig 4A**), progerin levels in M2I-1 treated HGPS cells were elevated when CQ was present, even though the two cell lines exhibited slight differences. We demonstrated in **Fig 3E** that progerin was in proximity to ubiquitin. Here, we tested whether progerin and ubiquitin were in proximity in HGADFN169 cells co-stained with antibodies against progerin and ubiquitin, following treatment with M2I-1. In the merged images of untreated cells, it can be seen that yellow foci are present within the nuclei, indicating that progerin and ubiquitin do indeed colocalize (**Fig 5C**). Similar to NB1-hTERT GFP-progerin cells (**Fig 4A**), M2I-I reduced progerin and ubiquitin levels in nuclei (**Fig 5C**). We observed that there was a change in the localization patterns of ubiquitin in M2I-1 treated cells, with low levels in the nucleus and cytoplasm and higher levels once HGPS cells were treated with M2I-1. This corresponds to a decrease in progerin pixel intensity within the nuclei of HGPS cells (**Fig 5D**). We surmise that activation of the APC by M2I-1 leads to increased ubiquitination of target proteins, including progerin. Taken together, these observations are consistent with the hypothesis that APC targets progerin for ubiquitin- and autophagy-dependent degradation (**Fig 6**).

## Discussion

Using computational tools, biochemical assays and imaging, we have identified a previously unknown interaction between the Anaphase Promoting Complex (APC) and the cytotoxic protein, progerin, the molecule responsible for the premature aging disease HGPS (21). We demonstrate the activation of the APC with the small chemical activator M2I-1 results in progerin degradation and a decrease in the prevalence of nuclear membrane blebbing. Evidence supports the hypothesis that progerin degradation resulting from APC activation occurs through ubiquitination-mediated autophagy.

Our first indication that there was potential that APC function may be affected by progerin expression in HGPS cells was observed in the meta-analysis of RNAseq data from cells isolated from HGPS patients (43). Multiple pathways are disrupted at the cellular level in HGPS patients, likely through the nuclear membrane defects or the loss of repressive epigenetic marks (62). Although there were other pathways that were disrupted in these datasets, the observation that APC subunits and targets are dysregulated (**Table S2-6**) suggested that the APC is acting aberrantly. The observation that APC function is altered in HGPS cells is parallel to what is observed in cancer cells (26, 42), where it was observed that reactivation of the APC lead to the death of drug resistant cancer cells (29, 41, 42, 63). This, and other evidence from yeast and cancer cell models (25, 26, 28, 29), support the role of the APC in impacting cellular aging and cancer progression.

We demonstrated that M2I-1 decreased progerin levels in a dose-dependent manner. We interpreted this to be due to APC activation based on previous published results (41, 42), as well as the decrease in levels of Cyclin B, a nAPC target (**Fig 1A**, **1B**). The APC mediates degradation of its targets through ubiquitination and targeting to the 26S proteasome (64). Surprisingly, we observed that progerin degradation is not inhibited by the proteasome inhibitor MG132 when the APC is activated (**Fig 1C-1E**), but Cyclin B degradation was blocked. This suggested that progerin may be targeted for degradation by the APC via autophagy. Indeed, it was previously shown that proteasome inhibition using MG132 induced auptophgy (21, 65). The APC itself has been shown to induce autophagy by targeting PFKKB3, an autophagy inhibitor, for degradation via the proteasome (66). However, there are no reports of the APC targeting substrates directly for autophagic breakdown. Furthermore, the decrease in progerin levels was associated with improvements in nuclear morphology and decreased doubling times (**Fig 2**). The increased doubling time suggests that growth is slowed to allow for improved cellular repair, similar to that reported for HGPS fibroblasts treated with Rapamycin (14).

If the APC does target progerin for ubiquitination, then the two proteins should be proximal to one another. Using Proximity Ligation Assays (PLA), we observed that progerin is in proximity to, and perhaps directly interacting with the APC (**Fig 3A-D**). Our PLA analyses indicate that progerin first interacts with CDC20 once CDC20 is free from the SAC. Next, we postulate that CDC20 then brings progerin to the APC to be ubiquinated (**Fig 6**). This is based on observations that treatment with APCIN, which inhibits the recruitment of CDC20 to the APC (67), blocks the proximal interaction of progerin with the APC2 subunit, but not the CDC20 co-activator (**Fig 3C**, **3D**). M2I-1 activates the APC by disrupting the interaction of CDC20 with the SAC (68). We observed increased association between CDC20 and progerin in the presence of M2I-1 (**Fig 3C**, **3D**), suggesting that CDC20 is inaccessible to progerin when bound to the SAC. Furthermore, APC activation using M2I-1 increased the interaction of progerin with ubiquitin, whereas APCIN blocked the interaction (**Fig 3E**). We further demonstrate that CDC20 co-immunoprecipitated with Lamin A/C antibodies (**Fig 3F**). We further demonstrate that progerin co-immunoprecipitated with CDC27 antibodies. Since CDC20 is recruited to the APC to activate it at the onset of anaphase once it is released from the SAC (35, 69), we propose that CDC20 carries progerin to the APC to trigger ubiquitin-mediated degradation (**Fig 6**).

Previous work has demonstrated that the ubiquitin ligase SMURF-2 targets progerin for degradation via autophagy (19). Several papers also present evidence that SMURF-2 may be involved with the APC. It has been shown that SMURF-2 may inhibit the APC by promoting SAC activity (70). Also, both the APC and SMURF-2 are recruited by Smad3 to target SnoN for ubiquitin-mediated degradation (71). Another study linked the Lamin A S143P mutation with decreased ubiquitin-proteasome system (UPS) activity, increased accumulation of Lamin A linked with K48-ubiquitin within the nucleus, and increased compensatory autophagic degradation of Lamin A-K48-Ub (72). There is therefore precedent for ubiquitination of progerin and other nuclear lamina associated proteins, indicating that there is a role for E3 ligases in mediating lamina synthesis and maintenance.

PLA and coimmunopreciptations provide evidence for interactions of the APC with progerin. Specifically, PLA demonstrates that APC2 is in proximity to progerin in HGPS cells and this is abolished in APCIN treated cells and that progerin increases its association with CDC20 in the presence of M2I-1. Further, PLA demonstrates that progerin is in proximity to ubiquin or other ubiquitinated proteins that increases in the presence of M2I-1. To support this, pulldown assays indicate that CDC27 is associated with progerin.

To provide further evidence that progerin is degraded by ubiquitin-dependant autophagy, we observed that chloroquine (CQ), an autophagy inhibitor, resulted in the accumulation of progerin in the presence of M2I-1 in NB1-hTERT GFP-progerin cells (**Fig 4A**). Although our data reveals that progerin is ubiquitinated in an APC-dependent manner, we cannot exclude that progerin is at least in proximity to ubiquitinated nuclear components that result in autophagy-mediated progerin degradation. If the APC is targeting progerin for autophagy, then M2I-1 should cause increased progerin ubiquitination that is transported out of the nucleus. However, when analyzed using immunofluorescence microscopy, we did not observe an increase in cytoplasmic GFP-progerin foci in CQ treated cells in the presence of M2I-1. We took advantage of the observation that GFP is resistant to autophagic destruction (61) to determine if M2I-1 does in fact increase free GFP. Within the cytoplasm of NB1 GFP-progerin cells, we observed low levels of GFP-progerin and that M2I-1 increased the amount of GFP-progerin and free GFP, but the levels did not reach significance (**Fig 4C, 4D**). CQ treatment significantly reduced free GFP in the cytoplasm, even in the presence of M2I-1, as expected for a chemical that inhibits autophagy. However, the combination of CQ and M2I-1 significantly increased the amount of GFP-progerin within the cytoplasm, indicating that M2I-1 does move progerin out of the nucleus to be targeted for autophagy. We also observed a shift from nuclear ubiquitin in control cells to elevated perinuclear ubiquitin with CQ + M2I-1 (**Fig 5D**), supporting the idea that M2I-1 does in fact move ubiquitinated nuclear proteins out of the nucleus.

Loss of protein homeostasis and protein accumulation is associated with aging and other age-related pathologies (73), in particular within the nuclear lamina. There is a significant body of research indicating the importance of protein homeostasis and the role of E3 Ub-ligases, such as the APC, in mediating this. Here, we reveal a previously unidentified interaction, either direct or indirect, between progerin and the activation if the APC with a small molecule inhibitor or MAD-2, results in progerin degradation and improved nuclear morphology. The schematic shown in **Fig 6** details the proposed pathway that the APC uses to target nuclear cytotoxic proteins for degradation. We do not have a clear understanding of how the APC triggers autophagy of cytotoxic proteins rather than canonical proteasome-dependent degradation of APC substrates. Future work will investigate the role of conserved autophagy pathway members in the targeting of proteins for APC-dependent autophagy. Our observations that CDC20 may interact with progerin prior to being recruited to the APC suggests a mechanistic difference with proteasome-dependent targeting of APC substrates. Our findings have significant implications into how we can take advantage of APC activation to further ameliorate the negative effects of cellular aging. Although there is significant hope in the near future for use of genetic modification or genome editing as a cure for HGPS, this will not be available for some time. While not a cure, our observations open a novel potential therapeutic avenue for the treatment of HGPS to ameliorate the effect of progerin expression.

## ACKNOWLEDGEMENTS

CHE and TAAH thank the Canadian Institutes for Health Research for funding this project. ZEG was supported by a scholarship from the Saskatchewan Innovation and Opportunity Fund and both ZEG and MF were supported by Vanier Canada Graduate Scholarships. ML was supported by a University of Saskatchewan College of Medicine Graduate Scholarship (CoMGRAD). CHE is also supported by the NSERC Discovery Grant Program.

## AUTHOR’S CONTRIBUTIONS

VM was responsible for all experiments using HGPS patient fibroblasts, PLA, and initial experiments requiring NB1-hTERT fibroblasts. ML performed cell fractionation and Westerns blots analyzing APC substrate protein levels. ML and MF performed immuno-precipitations. ZEG performed the meta-analysis of the RNAseq dataset from HGPS patients and controls. TAAH and CHE wrote the CIHR grant that funded the work. CHE supervised ZEG and VM, while TAAH supervised ML. VM and CHE contributed to the original writing. TAAH and CHE completed the writing and editing for submission for publication.

## FIGURE LEGENDS

**Supplemental Figure 1.** Principal Component analysis of HGPS patient and normal matched fibroblast RNAseq datasets. Three datasets from 3 male HGPS patients, 7 from female HGPS patients, and 18 from age-matched individuals were compared to show similarity of the datasets.

**Supplemental Figure 2.** APC2-CDC27 PLA interactions in HGADFN169 HGPS patient fibroblasts. HGADFN169 cells were grown in the presence of M2I-1, APCIN, or left untreated. Antibodies against APC2 and CDC27 were used to assess the relative proximity of the two proteins.

**Supplemental Figure 3.** APC2-progerin PLA interactions in HGADFN169 HGPS patient fibroblasts. HGADFN169 cells were grown in the presence of M2I-1, APCIN, or left untreated. Antibodies against APC2 and progerin were used to assess the relative proximity of the two proteins.

**Supplemental Figure 4.** NB1-hTERT normal fibroblasts were grown in culture, with antibodies against APC2 (A), CDC20 (B) and progerin used to assess the relative proximity of the proteins with progerin.

**Supplemental Figure 5.** HGADFN169 cells were used to determine the relative proximity of progerin and CDC20 in the presence or absence of M2I-1 or APCIN.

**Supplemental Figure 6.** HGADFN169 cells were used to determine the relative proximity of progerin and ubiquitin in the presence or absence of M2I-1 or APCIN.

**Supplemental Table 1.** List of SRA IDs for RNA-seq datasets used in meta-analysis (GEO: GSE113957), specifying age, biological sex, and disease status. “-” = unknown/undeclared

**Supplemental Table 2.** Differential Gene Expression (DESeq2) of HGPS fibroblasts (n=10) compared to age-matched controls (n=17) from Fleischer et al., 2018, GEO: GSE113957)

**Supplemental Table 3.** Genes identified as significantly increasing expression (absolute fold change ≥ 2, false discovery rate (FDR) < 0.05) by DESeq2 in HGPS fibroblasts compared to age-matched controls.

**Supplemental Table 4.** Genes identified as significantly decreasing expression (absolute fold change ≥ 2, false discovery rate (FDR) < 0.05) by DESeq2 in HGPS fibroblasts compared to age-matched controls.

**Supplemental Table 5.** Gene Ontology for all genes significantly changing expression (FDR < 0.05) between HGPS and age-matched controls

**Supplemental Table 6.** RNAseq dataset analysis reveals alterations in Anaphase Promoting Complex (APC) subunit and substrate mRNAs.

